# Pericyte control of pancreatic islet blood flow impacts glucose homeostasis

**DOI:** 10.1101/2021.08.24.457539

**Authors:** Alejandro Tamayo, Luciana Gonçalves, Rayner Rodriguez-Diaz, Elizabeth Pereira, Melissa Canales, Alejandro Caicedo, Joana Almaça

## Abstract

The pancreatic islet depends on blood supply to efficiently sense plasma glucose levels and deliver insulin and glucagon into the circulation. Long thought to be passive conduits of nutrients and hormones, islet capillaries were recently found to be densely covered with contractile pericytes, suggesting local control of blood flow. Here we determined the contribution of islet pericytes to the regulation of islet blood flow, plasma insulin and glucagon levels, and glycemia. Selective optogenetic activation of pericytes in intraocular islet grafts contracted capillaries and diminished blood flow. In awake mice, acute clamping of islet blood flow by optogenetic or pharmacological activation of pericytes disrupted hormonal responses, glycemia, and glucose tolerance. Our findings indicate that pericytes mediate vascular responses in the islet that are required for adequate hormone secretion and glucose homeostasis. Vascular deficiencies commonly seen in the islets of people with type 2 diabetes may impair regulation of islet blood flow and thus precipitate islet dysfunction.

## Introduction

Pancreatic islets play a crucial role in maintaining glucose homeostasis. There is general agreement that a functional islet vasculature is needed for adequate nutrient sensing, timely responses to changes in glycemia, and efficient release of insulin and glucagon into the circulation ^1,2^. Blood vessels in the pancreatic islet were long considered passive capillary tubes, but we recently established that islet capillaries are densely covered with contractile pericytes whose activity is tightly coupled to capillary constriction and dilation as well as to decreases and increases in blood flow in the islet [Fig. 1a; ^3^]. The density of pericytes is much higher in mouse and human islets than in the surrounding exocrine tissue of the pancreas and approximately half of pericytes in islet capillaries in mice and humans express high levels of the contractile protein alpha smooth muscle actin [Supplementary Fig. 1; ^3,4^], suggesting that pericytes work as internal gates (sphincters) that regulate blood flow at the level of the islet capillaries. Pericyte control of capillary blood flow, however, is still a source of debate. While some studies clearly showed that pericytes modulate capillary diameter and influence blood flow in tissues such as the brain, retina, and kidney ^5–9^, other studies reported that most of the control occurs upstream, at the level of pre-capillary arterioles ^10,11^. Addressing this issue may be particularly relevant for the pancreatic islet because such a local (capillary) control of blood flow may affect the secretion of glucoregulatory hormones that have systemic effects.

**Fig 1.**
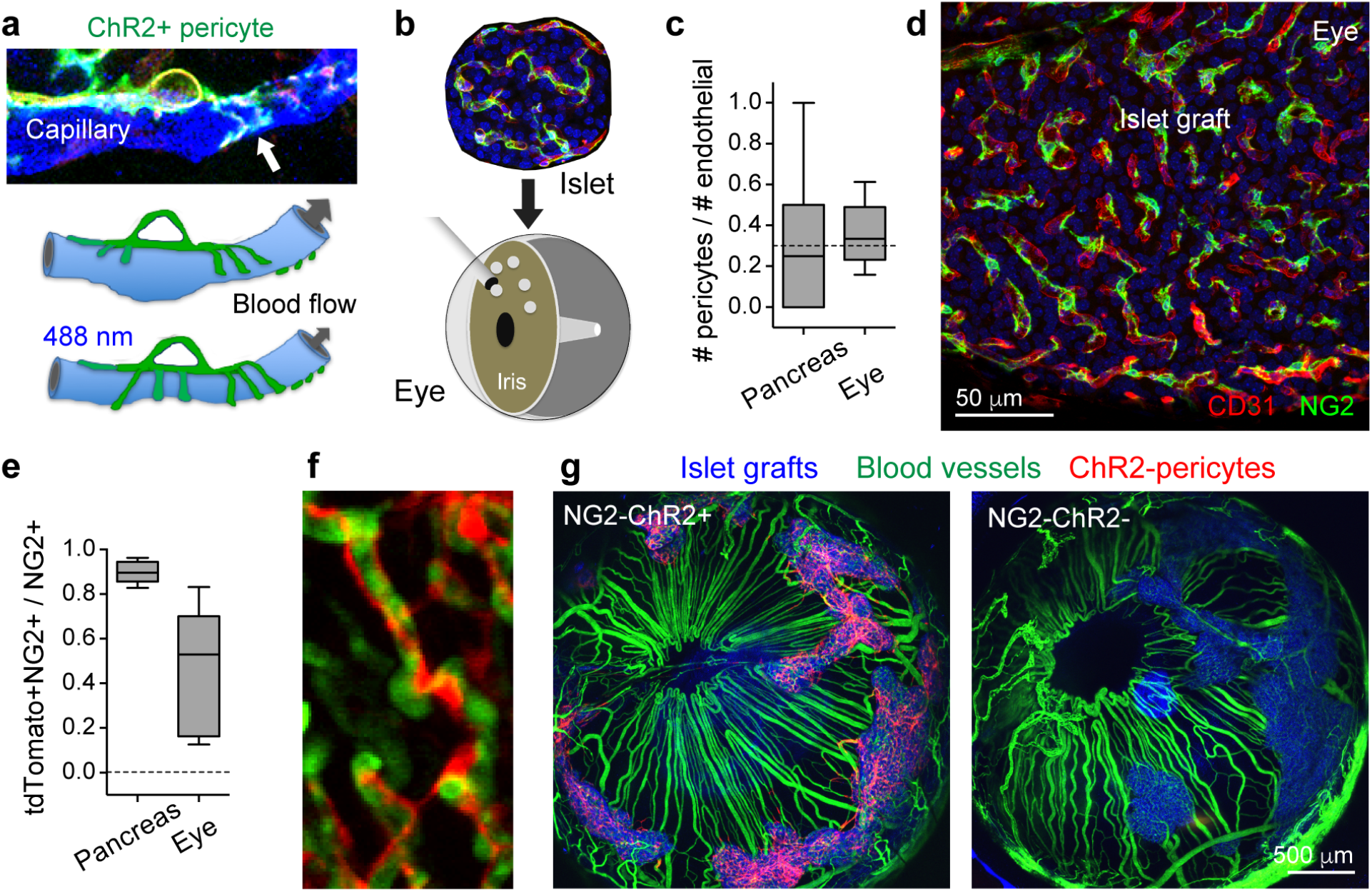
Experimental strategy to selectively activate islet pericytes. **a,b**, To visualize and manipulate islet pericytes, we generated mice that express channelrhodopsin (ChR2, fused to tdTomato) under the control of the NG2 promoter (NG2-ChR2+ mice and control NG2-ChR2- mice) and transplanted 200 islets from these mice into the eyes of wildtype mice rendered diabetic with streptozotocin. **c**, Pericyte coverage (ratio pericytes to endothelial cells) of capillaries in islets in the pancreas and in intraocular grafts was similar. **d**, Z-projection showing pericytes (NG2, green) covering capillaries (CD31, red) in an intraocular islet graft. **e**, Mander’s coefficient reflecting ChR2 (tdTomato) and NG2 colocalization for islet pericytes in the pancreas and the eye. **f**, *In vivo* imaging shows that pericytes (red) remain associated with islet capillaries in the islet graft (FITC dextran, iv). **g**, Islets (backscatter signal in blue) fully engraft and control glycemia one month after transplantation into the eyes of diabetic mice. In mice transplanted with islets from NG2-ChR2+ mice, but not from NG2-ChR2- mice, pericytes can be selectively stimulated with blue light because they express ChR2 (red, left).

In previous studies, we had determined that pericyte function is regulated by local signals derived from insulin secreting beta cells (e.g. adenosine) and sympathetic nerves. Importantly, along with other vascular abnormalities of the islet, pericytes progressively disappear in diabetes ^3^, but the pathophysiological consequences remain unknown. Unfortunately, our understanding of the importance of islet blood flow for glucose homeostasis has depended on inferences from indirect studies based on manipulation of the systemic vasculature and circulation ^12–14^, or on chronic ablation of islet blood vessels ^15–17^. Here we investigated how islet blood flow directly impacts pancreatic islet function and systemic glucose metabolism.

### Selective manipulation of pancreatic islet blood flow using optogenetics and transplantation into the eye

To test the hypothesis that acute control of islet blood flow affects glucose metabolism, we used an experimental approach that allowed direct and selective manipulation of islet pericytes. We transplanted islets into the anterior chamber of the eye of mice rendered diabetic with the beta cell toxin streptozotocin (Fig. 1b). This model has been used extensively to study islet blood vessels because the revascularization of the graft faithfully reconstitutes the structural elements of the vasculature of the islet in the native pancreas ^18–26^. Of note, intraocular islet grafts restore and maintain normoglycemia a couple of weeks after transplantation (Supplementary Fig. 2), allowing to correlate local manipulation of islet cells to systemic effects.

To manipulate pericytes, we transplanted islets isolated from mouse donors expressing the light-gated ion channel channelrhodopsin under the control of the pericyte promoter NG2 (NG2-ChR2+ mice; Figs. 1a,b; Supplementary Fig. 3). Donor pericytes not only survived the islet isolation and transplantation procedures, but also covered the islet endothelial network as they do in the pancreas (Figs. 1c,d) without colonizing the surrounding vasculature (Fig. 1g and Supplementary Fig. 4b). Pericytes in islet grafts from NG2-ChR2+ donor mice, but not NG2-ChR2- mice, expressed channelrhodopsin (Figs. 1e-g and Supplementary Fig. 4c,d). Once islets fully engrafted on the iris and took over the control of glucose metabolism, we manipulated pericytes optogenetically or pharmacologically and determined the effects on capillary diameter, islet blood flow, plasma hormone levels, and glycemia.

### Pericytes control islet blood flow and glucose diffusion

To determine how activation of pericytes affects capillaries, we first stimulated intraocular islet grafts with a 488 nm laser while imaging capillary diameter and blood flow under a confocal microscope. Activating pericytes with blue light diminished the diameter of ~30% of islet capillaries (32.5 ± 10.4%; Figs. 2a,b; Supplementary Movie 1). Changes in capillary diameter ranged from 20 to 50% reduction of the initial value. We tested the impact of pericyte activation on islet blood flow and found that blood flow dropped significantly in ~80% of islet capillaries (Figs. 2c-g). In some capillaries (70% of responsive vessels), pericyte constriction led to a complete stop of blood flow (Figs. 2f,g; Supplementary Fig. 5; Supplementary Movies 2-4). Next, we investigated if light stimulation of pericytes limited plasma glucose diffusion into the islet parenchyma. We injected fluorescent glucose into the circulation and found that activating pericytes diminished glucose diffusion into the islet parenchyma (Figs. 2h,i). Thus, optogenetic activation of islet pericytes contracted capillaries, decreased blood flow, and limited perfusion of the islet. We could not observe any vascular effects when we used red instead of blue light to stimulate or when pericytes did not express channelrhodopsin (Supplementary Fig. 5).

**Fig 2.**
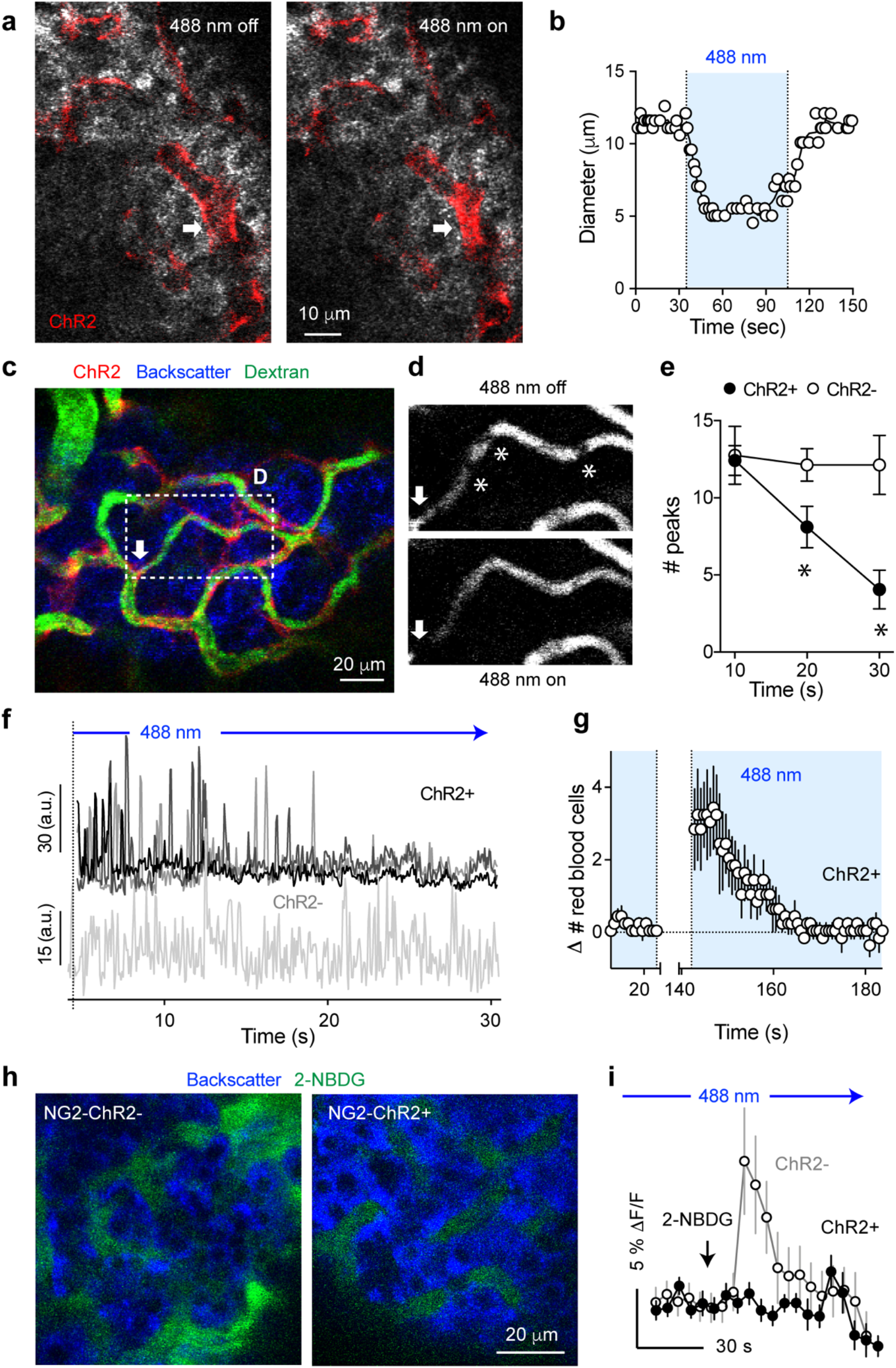
Optogenetic activation of pericytes constricts islet capillaries, decreases blood flow, and limits perfusion of the islet. **a,b**, Confocal images taken during *in vivo* imaging of an intraocular islet graft show that activating ChR2-expressing pericytes with blue light (488 nm laser) constricted an islet capillary (arrow; changes in diameter are plotted in **b**). **c**, *In vivo* confocal image of an islet graft showing islet vessels (dextran, green) and ChR2-expressing pericytes (red). Arrow indicates a potential “sphincter” pericyte. **d**, Blood flow through an islet graft capillary (region shown in (**c**)) is reduced upon turning on the 488 nm laser. * indicate different erythrocytes (shadows). **e**, Quantification of the number of peaks in the first 10, 20 or 30 sec after turning on the 488 nm laser, from traces as in (**f**) N = 9-19 vessels, from 3 NG2-ChR2- (white) and 3 NG2-ChR2+ (black); Supplementary Fig. 5]. **f**, Traces showing changes in mean fluorescence in ROIs placed on different capillaries in islet grafts from control NG2-ChR2- (bottom panel) and NG2-ChR+ mice (top panel) after performing Stack Difference (see Methods). **g**, Quantification of blue light-induced changes in the density of erythrocytes in islet capillaries as shown in **d** (n = 5 vessels). 488 nm laser had been on for 1 min before starting *in vivo* imaging. **h**, Diffusion of the fluorescent glucose analog 2-NBDG (5 mg/kg, iv) was reduced by optogenetic stimulation started at 30 s before glucose injection. *In vivo* images of 2-NBDG fluorescence (green) in islet grafts from control NG2-ChR2- and NG2-ChR2+ mice 30 s after injection. **i**, Quantification of changes in 2-NBDG fluorescence intensity as shown in **h** (intensity levels in the islet graft parenchyma were normalized to those in the aqueous humor; arrow indicates time of injection).

### Pericytic control of islet blood flow impacts glucose homeostasis

To investigate how manipulating control of blood flow in the islet impacts systemic glucose metabolism, we performed studies in awake, freely moving, transplanted mice (Fig. 3a). After 4 hours of light exposure in cages equipped with blue LED light for ChR2 activation (1 min on, 4 min off), hypoxic levels and nitric oxide production increased in islet grafts, confirming that the manipulation affected vascular perfusion of islet grafts (Figs. 3b-d). We then tested how acute light stimulation (10 min) affected islet hormone secretion and glycemia. In control mice transplanted with islets from NG2-ChR2- donors, light elicited changes in plasma insulin and glucagon levels (Fig. 3e), most likely due to autonomic input to islet grafts in the context of the pupillary light reflex ^27,28^. In mice transplanted with islets from NG2-ChR2+ donors, this hormonal response was reversed, indicating that activating light sensitive pericytes affected hormone release into the circulation (Figs. 3e-g). This disrupted hormone secretion changed the insulin to glucagon ratio and destabilized glycemia (Supplementary Fig. 6).

**Fig. 3.**
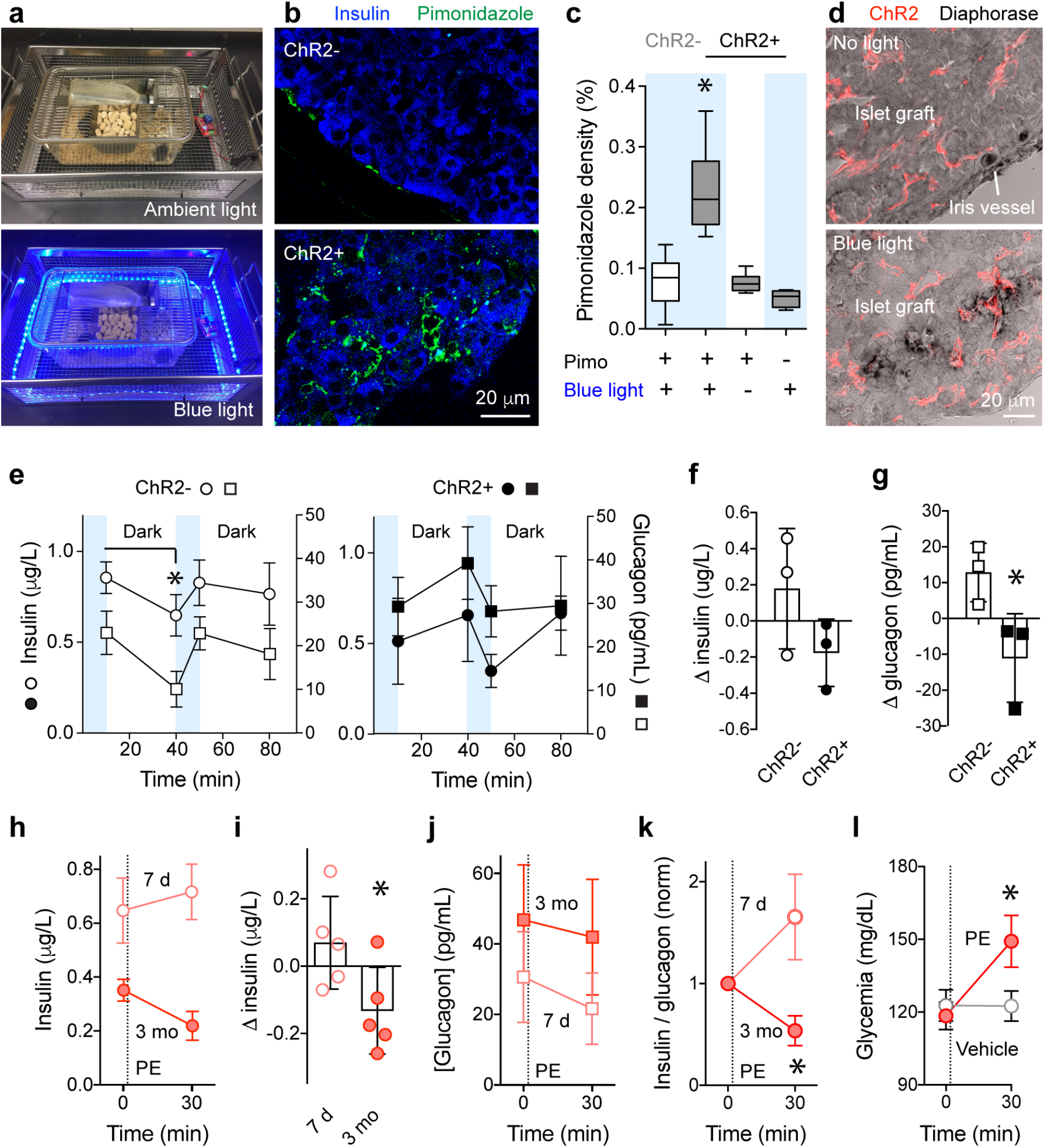
Islet pericyte activation impacts hormone secretion and glycemia in awake, freely moving transplanted mice. **a**, Mice with intraocular islet grafts from NG2-ChR2+ and NG2-ChR2- mice were housed together in cages equipped with a blue LED light around their perimeter and subjected to the same light stimulation protocol. **b**, Stimulation with blue light pulses for 4 h (1 min on, 4 min off) increased levels of the hypoxia marker pimonidazole (60 mg/kg, green) in mice with NG2-ChR2+ islets (bottom) but not in mice with NG2-ChR2- islets (top). **c**, Quantification of results as in **b** showing the density of pimonidazole immunostaining (immunostained area / graft area) in islet grafts (**c,** *p<0.05, one-way ANOVA Tukey’s multiple comparisons test, n = 4-6 grafts/6 mice). **d**, Optogenetic activation of islet pericytes increased NADPH diaphorase activity (black staining) in grafts from NG2-ChR2+ mice (bottom) but not in the absence of light (top). Notice ChR2-tdTomato expressing pericytes in red and background NADPH diaphorase activity in iris vessels. **e**, Acute exposure to blue light (10 min) increased plasma insulin (circles) and glucagon (squares) secretion in control mice (NG2-ChR2-; white symbols; left panel) but decreased both hormones in mice with ChR2 expressing islet pericytes (NG2-ChR2+; black symbols; right panel). **f,g**, Quantification of results presented in **e**. Shown is the difference (Δ) between hormone levels at the beginning and end of the 10 min period. (*p < 0.05, unpaired Student’s t-test, n = 3 mice). **h-l**, Eyes of C57BL6 mice transplanted with syngeneic islets were treated with topical application of the a1 adrenergic receptor agonist phenylephrine (PE; eye drops, 2.5%) at 7 days (7 d; pink symbols) and 3 months (3 mo; red symbols) after transplantation, and plasma insulin (**h** and **i**) and glucagon (**j**), their ratios (**k**), and glycemia (**l**) were determined. The PE-induced change in plasma insulin (**I**, *p < 0.05, paired t-test, n = 5 mice), decrease in the insulin / glucagon ratio (**k**, *p < 0.05, paired t-test, n = 5 mice), and increase in glycemia (**l**, *p < 0.05, One-way ANOVA Sidak’s multiple comparisons test, n = 5 mice) were significantly different at 3 months. Ratios were normalized to values before PE.

To determine how vascular manipulation affects hormone secretion and glycemia independently of light stimulation, we used a pharmacological approach to target islet pericytes. Applying phenylephrine, an alpha 1 adrenergic receptor agonist, did not affect islet hormone secretion *in vitro* but elicited responses in pericytes, constricted capillaries, and reduced blood flow in intraocular islet grafts [Supplementary Fig. 7; ^3^]. Thirty minutes after topical application of phenylephrine (eye drops, 2.5%), plasma insulin levels decreased in mice with fully engrafted islets (3 months after transplantation), but not in non-transplanted mice or in the same mice early after transplantation (7 days), when transplanted islets have not been revascularized yet (Figs. 3h,i). Although plasma glucagon levels did not change, the insulin to glucagon ratio decreased and glycemia increased (Figs. 3j-l).

The optogenetic and pharmacological studies indicated that preventing local control of islet blood flow affects islet hormone secretion and glycemia. We next addressed the role of islet blood flow in the islet’s response to a glycemic challenge. We conducted intraperitoneal glucose tolerance tests (IPGTTs) in awake, freely moving, transplanted mice. Activating pericytes in intraocular islet grafts of mice placed in cages equipped with blue light increased the glucose excursion and made mice glucose intolerant (Figs. 4a-c). Red light did not affect glucose tolerance (data not shown). Mice with control islets from NG2-ChR2- donors showed the typical hormone response to a glucose challenge, that is, an increase in insulin and a decrease in glucagon levels (Fig. 4d). By contrast, mice with activatable pericytes in their islet grafts from NG2-ChR2+ donors showed a reversed hormone response. Insulin secretion was reduced, and glucagon secretion increased (Fig. 4d), thus abolishing the expected change in the insulin to glucagon ratio under these conditions (Fig. 4e).

**Fig. 4.**
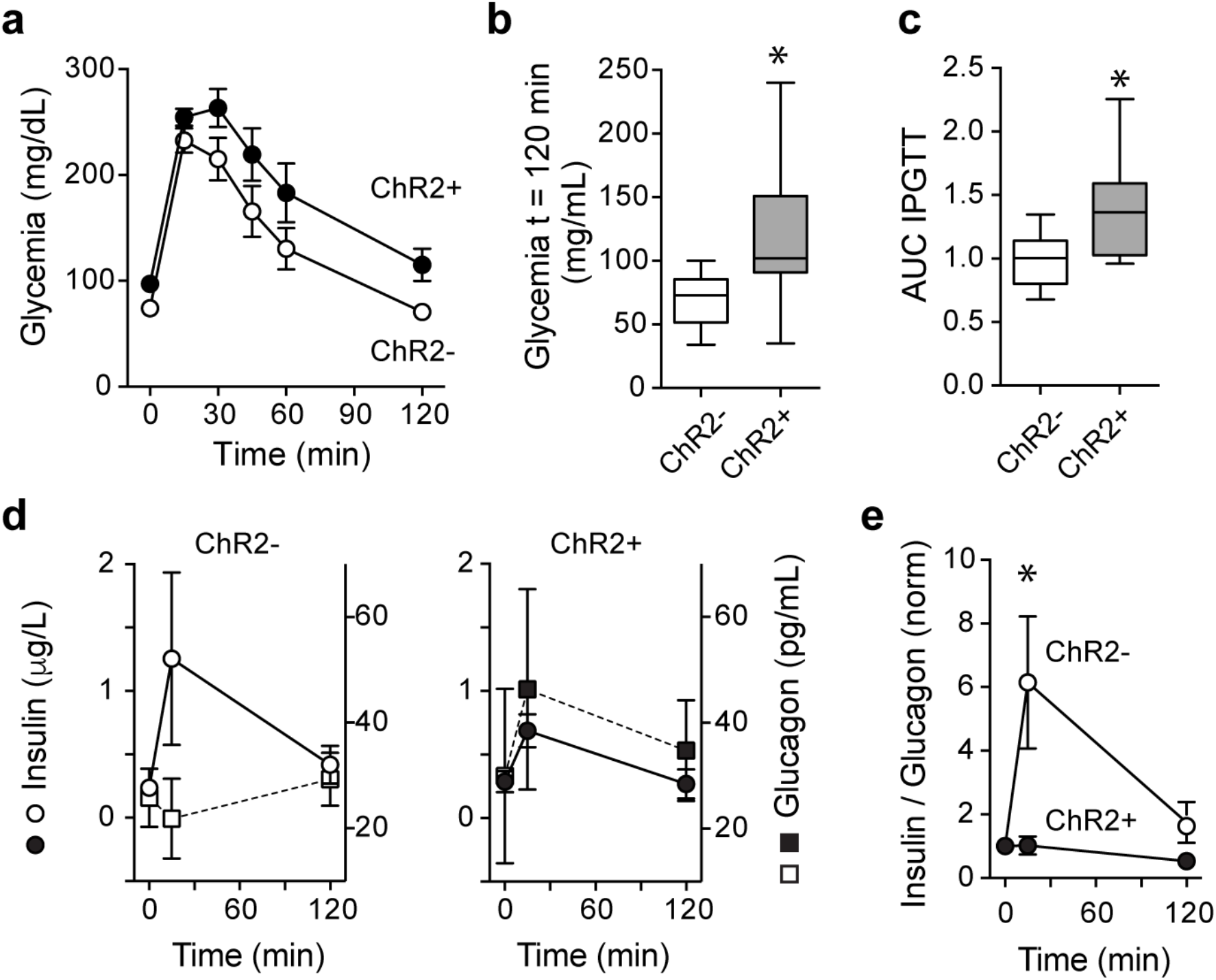
Optogenetic activation of islet pericytes impairs the islet response to a glucose challenge. **a**, Glucose excursions of an intraperitoneal glucose tolerance test (IPGTT) performed in awake, freely moving mice transplanted with islets from NG2-ChR2+ (black) and NG2-ChR2- mice (white) in the presence of blue light pulses (1 min on, 4 min off; light pulses started 1h before and continued during the IPGTT; n = 7-9 mice/group). **b**, Glycemia at 120 min after glucose injection. **c**, Area under the curve (AUC) of IPGTTs performed upon stimulation with blue light (*p < 0.05, unpaired t-test, n = 7-9 mice/group). Values were normalized for AUC values from NG2-ChR2- mice. **d**, Changes in plasma insulin (circle) and glucagon (square) induced by glucose in control mice (white symbols; left panel) and in mice with ChR2 in islet pericytes (black symbols; right panel) during the IPGTT. **e**, Optogenetic activation of islet pericytes abolished the increase in the insulin / glucagon ratio induced by the glucose challenge (*p < 0.05, multiple t-tests, n= 5-6 mice). Ratios were normalized to values before glucose administration.

## Conclusions

Our results argue against the idea that islet capillaries are passive conduits for nutrients and hormones. On the contrary, we show that the islet vasculature is endowed with pericytes that actively modulate local blood flow, thus affecting hormone secretion from the islet and impacting glycemic control. We had previously reported that islet pericytes respond to signals derived from insulin secreting beta cells and sympathetic nerves ^3^. Therefore, our findings indicate that islet pericytes couple endocrine and neural inputs to vascular responses needed for proper context-dependent hormone secretion. The pericyte may mediate the reported functional hyperemia of the islet ^29–34^, enabling adequate hormonal responses under hyperglycemic conditions (Fig. 4). By responding to sympathetic input, pericytes may also contribute to the islet response that counters hypoglycemia (Fig. 3). This neurovascular coupling may be particularly relevant for human islet function. Autonomic nerves predominantly target the islet vasculature [Supplementary Fig. 7, ^35^] and activation of sympathetic nerves in the pancreas has been shown to decrease plasma insulin levels in human beings ^36^.

Vascular abnormalities such as loss of pericytes, amyloid accumulation, and fibrosis are commonly present in the islet of people with type 2 diabetes ^37^. These vascular defects may disrupt endo- and neuro-vascular coupling and chronically impair local regulation of blood flow, which may contribute to diabetes pathogenesis.

## Methods

### Transgenic mice

Mouse lines used in this study were from the Jackson Laboratory and Envigo (USA). For optogenetic manipulation, we crossed male mice that have floxed expression of the blue light sensitive cation channel Channelrhodopsin2 (fused to tdTomato fluorescent protein; ChR2-tdTomato-floxed, JAX stock no. 012567) with female mice that express Cre recombinase under the *Cspg4* promoter/enhancer (NG2-Cre, JAX stock no. 008533). Because NG2-Cre mice are kept as heterozygous, the breeding scheme used produced mice that have ChR2 in pericytes (Cre+; NG2-ChR2+; experimental) and control mice that do not have ChR2 in pericytes (Cre-; NG2-ChR2-; control). Glucose metabolism, islet endocrine cell composition and vascular architecture is similar in NG2-ChR2+ and NG2-ChR2- mice (Supplementary Fig. 3). Islets from NG2-ChR2+ and NG2-ChR2- female and male mice (4-6 months of age) were isolated and transplanted into the anterior chamber of the eye of immune-compromised, athymic nude mice (Hsd:Athymic Nude-nu; age: 8 weeks; Envigo). Male and female Nude mice were used as transplantation recipients and yielded similar results (Supplementary Fig. 2). For pharmacological manipulation of pericytes *ex vivo*, we crossed male mice that have floxed expression of GCaMP3 (GCaMP3-floxed, JAX stock no 029043) with female NG2-Cre mice. For pharmacological manipulation of pericytes and islet blood flow *in vivo*, we used C57BL6 male and female mice (JAX stock no. 000664; age 12 weeks) as islet donors and recipients. All animals were housed in virus antibody-free (VAF) rooms and kept in micro-isolated cages (5 mice per cage) with free access to autoclaved food and water. All experiments were conducted according to protocols approved by the University of Miami Institutional Animal Care and Use Committee.

### Islet isolation and *in vitro* hormone secretion

Pancreatic islets were isolated by cannulation of the common bile duct of donor animals and infusion of 3 ml collagenase P, 1.0 mg/ml in HBSS containing 25 mM HEPES (Sigma-Aldrich, USA). Inflated pancreata were dissected out and digested at 37°C for 15 min. Islets were cultured in CMRL Medium (5.5 mM D-glucose) supplemented with 10% fetal bovine serum, 100 IU/ml penicillin, 100 μg/ml streptomycin and 2 mM L-Glutamine (all from Thermo Fisher Scientific, USA). Perifusion studies of isolated islets was done as previously described ^22^.

### Transplantation into the anterior chamber of the eye

Recipient mice were rendered diabetic before islet transplantation with a single intravenous injection of the beta cell toxin streptozotocin (STZ 200 mg/Kg; Sigma, MO) dissolved fresh in 100 mM sodium citrate (pH 4.5) immediately before injection. There was no morbidity or mortality associated with streptozotocin toxicity.

Transplantation into the anterior chamber of the eye was performed as previously described ^22,25^. Eyes were kept humidified (ophthalmologic eye drops) to avoid drying of the cornea. Under a stereomicroscope, the cornea was punctured close to the sclera at the bottom part of the eye with a 31G insulin needle. Using the needle, we made a small radial incision of approximately the size of the eye cannula (~ 0.5 mm). For this incision, the needle was barely introduced into the anterior chamber, thus avoiding damage to the iris and bleeding. The blunt eye cannula was then gently inserted through this incision, first perpendicular to the surface of the cornea and then parallel to the cornea. Once the cannula was stably inserted into the eye, the islets were slowly injected in a 10-μl volume of sterile saline solution into the anterior chamber, where they settle on the iris. After injection, the cannula was carefully and slowly withdrawn (1 min) to avoid islets from flowing back through the incision. The mouse was left lying on the side before awakening. Mice were then put back in the cages and monitored until full recovery. Analgesia was obtained after surgical procedures with buprenorphine (0.05-0.1 mg/kg s.c.).

Only animals with non-fasting glycemic values >450 mg/dl were transplanted. We transplanted 200 mouse islet equivalents from NG2-ChR2+ and NG2-ChR2- mice into the anterior chamber of the right eye of diabetic Nude recipient mice (for optogenetic manipulation) and from C57BL6 mice into diabetic B6 siblings (for pharmacological manipulation with phenylephrine eye drops). This provided an optimal beta cell mass that allowed reversal of diabetes for 26 mice out of 28 recipient mice (total number of transplanted mice for both manipulations) within 1 month after transplantation. The 2 animals that never recovered from diabetes received NG2-ChR2+ islets (see Supplementary Fig. 2) and were not included in the study. We defined normoglycemia as 3 consecutive readings of non-fasting blood glucose below 200 mg/dl (2, 4, 5).

### *In vivo* imaging of the mouse eye

Imaging of islets *in vivo* in the anterior chamber of transplanted animals was performed as previously reported. Briefly, mice were anesthetized with ~2% isoflurane air mixture, placed on a heating pad and the head restrained with a headholder. The eyelid was carefully pulled back and the eye gently supported. For fluorescence confocal imaging, an upright SP5 Leica confocal microscope (Leica Microsystems, Mannheim, Germany) was used for imaging together with long distance water-dipping lenses (Leica HXC APO 20x 0.5 W), using Viscotears (Novartis, Basel, Switzerland) as an immersion liquid. ChR2-tdTomato expressing pericytes were visualized upon excitation with 561 nm laser and reflected light was imaged by illumination at 633 nm and collection between 630-639 nm. To activate/deactivate ChR2, we turned on/off the 488 nm laser during image acquisition (100% laser power). Blood vessels were labeled by tail vein injection of 150,000 Da Dextran-FITC. FITC was excited at 488 nm and emission light was collected between 500-550 nm. For monitoring the uptake of glucose analog, 2-(N-(7-Nitrobenz-2-oxa-1,3-diazol-4-yl)Amino)-2-Deoxyglucose (2-NBDG, Thermo Fisher Scientific, USA) was dissolved in saline solution at 5 mg/ml. 100 μl 2-NBDG solution was injected i.v. Time series of z-stacks (2 μm) were acquired every 5 sec.

Data analysis was performed with ImageJ (http://imagej.nih.gov/ij/; NIH, USA) software. The density of FITC-labeled blood vessels in each islet graft was quantified by calculating the area of stained blood vessels as a fraction of the islet area (vessel density). Changes in capillary diameter induced by optogenetic manipulation of pericytes were determined as previously described ^3^. Briefly, we drew a straight-line transversal to the blood vessel borders and used the “reslice” z-function in ImageJ to generate a temporal projection ^38^. We drew another line on the temporal projection and, using the “plot profile” function, we determined the position of the pixels with the highest fluorescence intensity and considered these the vessel borders. Vessel diameter was calculated by subtracting these 2 position values. To analyze changes in blood flow pattern, after stabilizing our *in vivo* recordings using the ImageJ plugin “Image Stabilizer”, we used the “Stack Difference” plugin to detect dynamic pixel changes ^39^. We drew regions of interest (size of an individual erythrocyte) on different vessels and quantified changes in fluorescence over time (Supplementary Fig. 5 and Movie 3). By simultaneous inspection of our *in vivo* recordings and corresponding traces of changes in fluorescence, we counted only peaks whose maxima corresponded to a change in fluorescence greater than 40% of the baseline. Peaks in traces were manually counted in intervals of 10s (10, 20 and 30 s bins were used) after turning on the 488 nm laser.

### Optogenetic manipulation and assessment of metabolic function

We monitored graft function in transplanted animals by measuring non-fasting glycemia using a portable glucometer (OneTouch, LifeScan) and body weight. Tail blood samples (20 μL) were taken for determination of plasma insulin and glucagon levels in the nonfasting state at different time points after transplantation in the absence of blue light stimulation (Supplementary Fig. 2). Manipulation with blue light to activate ChR2 started 3 months after transplantation when all the animals were normoglycemic.

To activate ChR2 expressed by pericytes in intraocular islet grafts, twenty blue light-emitting diodes (LEDs, emission peak at 465 nm, Cree, Durham, NC, USA) were fixed on a special ion cage and the cages housing transplanted mice were placed in the ion cage, as previously described [^40^; (Fig. 3a)]. As a control, cages with red LEDs were also built. The output power of each LED is 10 mW. Blue light was delivered chronically for 10 min (Figs. 3, e-g) or in pulses (rest of the manuscript figures). The frequency of pulsed blue light was controlled by a digital cycle timer switch that can be programmed to have different frequency settings (Inkbird, Shenzhen, China). The blue light was set to keep off for 4 min and on for 1 min. Mice transplanted with islets from NG2-ChR2+ (Cre+; experimental) or from NG2-ChR2- (Cre-; controls) were always housed together and subjected to the same light stimulation protocol. Stimulation with blue light occurred only acutely, and otherwise transplanted mice were kept in control environment in normal 12 h light/dark cycle.

To assess changes in glucose tolerance, mice transplanted with NG2-ChR2+ or NG2-ChR2- islets were placed in the dark and fasted overnight to perform an intraperitoneal (i.p.) glucose tolerance test (IPGTT). Stimulation with pulsed blue light started 1h before the IPGTT and continued throughout the test. We measured glycemia following i.p. injection of glucose (2 g/kg). As a control, we performed an IPGTT with the same mice in the presence of pulsed red light. To determine changes in glucose-stimulated hormone secretion, transplanted mice were fasted for 6h. Stimulation with pulsed blue light started 4h before glucose injection and continued throughout the test. Blood was collected from the tail vein before, 15 min and 120 min after glucose injection. Plasma insulin and glucagon levels were determined using an ultra-sensitive mouse insulin or a glucagon Elisa Kit following manufacturer’s instructions (Mercodia).

### Assessment of tissue hypoxia and nitric oxide production

Transplanted mice were placed in the dark and stimulated with pulsed blue light (4 min off, 1 min on) for 4h. The oxygenation marker pimonidazole (60 mg/Kg; dissolved in sterile saline solution; Hypoxyprobe™ kit) was injected intravenously and allowed to circulate for 90 min. Pimonidazole that precipitates at oxygen levels of ≤8 mmHg ^15^. Control animals included mice with intraocular NG2-ChR2- islet grafts stimulated with blue light and treated with pimonidazole, and mice with NG2-ChR2+ islet grafts either not stimulated with pulsed blue light or treated with saline solution. After 90 min, animals were sacrificed and the eyes with islet grafts were dissected, fixed in 4% paraformaldehyde (overnight) and processed for immunohistochemistry.

To assess local nitric oxide levels, we used NADPH-diaphorase histochemistry technique. NADPH diaphorase is commonly used as a histochemical marker of nitric oxide synthase and other nitric oxide-containing factors in aldehyde-fixed tissues ^41^. Briefly, frozen sections of the eyes containing grafts (10 μm sections) were rinsed twice in 50 mM Tris-buffer (pH 8.0) and incubated for 45 min in Tris-buffer containing 0.5% Triton X-100, before incubating with nitroblue (0.2 mM) and NADPH (1 mM) at 40’C for 15-30 min. Sections were rinsed, mounted and imaged on a confocal microscope.

### Pharmacological activation of pericytes in intraocular islet grafts

As an alternative manipulation of pericyte activity independently of light, we transplanted diabetic C57BL6 mice with islets from sibling B6 mice. Early after transplantation (7 days), when islets had still not engrafted (fed glycemia > 300 mg/dL) and when fully revascularized (3 months after transplantation; fed glycemia < 200 mg/dL), transplanted animals were fasted for 6h and treated with phenylephrine containing eye drops (2.5%, one drop in transplanted eye). Glycemia and blood were collected before and 30 min after PE drop. A group of mice was treated with vehicle. To confirm that phenylephrine activates pericytes, we produced living pancreas slices from NG2-GCaMP3 mice as previously described ^3^ and recorded changes in cytosolic calcium levels induced by phenylephrine (10 and 100 μM) *ex vivo*.

### Immunohistochemistry

Mice expressing ChR2 in pericytes and mice with intraocular NG2-ChR2 islet grafts were perfused with 4% PFA and the pancreases and eyes dissected, fixed overnight in 4% PFA, cryoprotected in a sucrose gradient (10, 20 and 30% w/w sucrose), and frozen in Tissue-Tek Optimal Cutting Temperature (OCT) compound before cryosectioning (−20°C). After a rinse with PBS-Triton X-100 (0.3%), sections were incubated in blocking solution (PBS-Triton X-100 and Universal Blocker Reagent; Biogenex, San Ramon, CA). Thereafter, sections were incubated 48 hr with primary antibodies diluted in blocking solution. We immunostained beta cells (insulin; Accurate Chemical & Scientific, Wesbury, NY), alpha cells (glucagon; Sigma, St. Louis, MO), delta cells (somatostatin), endothelial cells (PECAM; BD Biosciences, San Jose, CA), and pericytes (neuron-glial antigen 2; NG2; Millipore). tdTomato was used as a reporter of ChR2 expression and amplified with an antibody against mCherry. Immunostaining was visualized by using Alexa Fluor conjugated secondary antibodies (1:500 in PBS; 16 hr; Invitrogen, Carlsbad, CA). Pimonidazole is reductively activated in hypoxic cells and forms stable adducts that can be detected with a fluorescent antibody (FITC-Mab1; Hypoxyprobe™). Cell nuclei were stained with DAPI. Slides were mounted with Vectashield mounting medium (Vector Laboratories) and imaged on a confocal microscope. ImageJ was used to quantify endocrine and vascular cell numbers in confocal planes. To determine colocalization between NG2 and ChR2 (tdTomato), we used the ImageJ plugin “intensity correlation analysis” and calculated the Mander’s Colocalization coefficients (M1 and M2). These-coefficients avoid issues related to the absolute intensities of the signals, since they are normalized against total pixel intensity ^42^, and represent the fraction of NG2 that is present in ChR2/tdTomato positive structures (M1) and vice – versa (M2). Quantifications were performed in confocal images, in a minimum of 3 islet/grafts per mice/3 mice per group.

### Statistical analysis

Throughout the manuscript, the results are expressed as mean ± S.E.M or as otherwise specified from at least three independent biological experiments. A value of *P* < 0.05 was considered statistically significant. All statistical tests, sample sizes and their *P* values are provided in the figure legends. All statistics were performed using GraphPad Prism Software.

## Acknowledgements

We would like to thank Drs. Vlad Slepak, Steve Roper, Ellen Barrett and John Barrett for carefully reviewing the manuscript. This work was funded by NIH grants [K01DK111757 (J.A.), R01DK08432, R01DK111538, R01DK113093 (A.C.) and by the NIDDK-supported Human Islet Research Network (HIRN, RRID:SCR_014393; https://hirnetwork.org; UC4 DK104162, New Investigator Pilot Award to J.A.).

## Author contributions

A.T., L.M.G, E.P. and J.A. performed experiments; R.R.D performed islet transplantation into the anterior chamber of the eye; M.C. and J.A. analyzed data; J.A. and A.C. designed the study and wrote the paper. All authors discussed the results and commented on the manuscript.

## Competing interests

All authors declare that they have no competing interests.

## Data and materials availability

All data are available in the main text or the supplementary materials. Requests should be sent to Joana Almaça.

## Notes

### Competing Interest Statement

The authors have declared no competing interest.

