## Supplementary Figure for "Pericyte control of pancreatic islet blood flow impacts glucose homeostasis"

### Supplementary Material

### Supplementary Figures

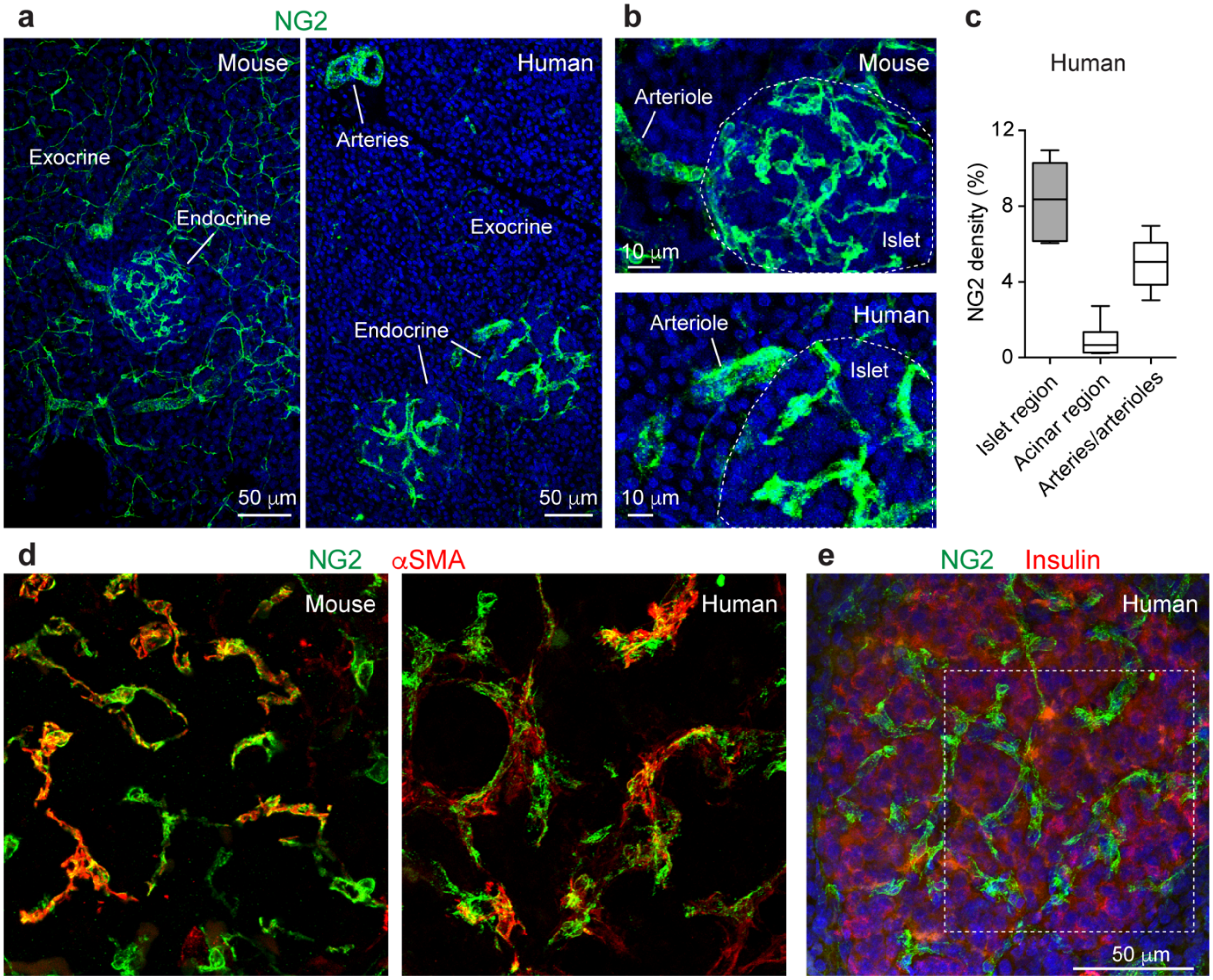


**Supplementary Fig. 1. Pancreatic islets are full of contractile pericytes. a**, Z-projection of confocal images of mouse and human pancreatic tissue sections immunostained with an antibody against the mural marker NG2 (neuron-glial antigen 2; green). In humans, besides smooth muscle cells around arteries or arterioles, NG2-positive cells are only found in islets. **b**, Pericyte density in mouse and human islets is high. **c**, Quantification of NG2 immunostained area in regions in the human pancreas (of similar area) containing islets, acinar tissue or larger blood vessels such as arterioles or arteries. **d**, Z-projection of confocal images of a mouse and human pancreatic tissue section immunostained for pericytes (NG2, green) and alpha smooth muscle actin

($\alpha$SMA, red). Around 50% of mouse and human islet pericytes express $\alpha$SMA as previously reported [1,2]. **e**, Regions rich in pericytes in the human pancreas are islets. Insulin-positive beta cells are shown in red and pericytes in green. Zoomed image of region within square is shown in **d**. These data suggest that pancreatic islets in mice and humans have a high density of contractile pericytes, being thus equipped with a mechanism that allows the control of their blood flow independently of the surrounding exocrine tissue.

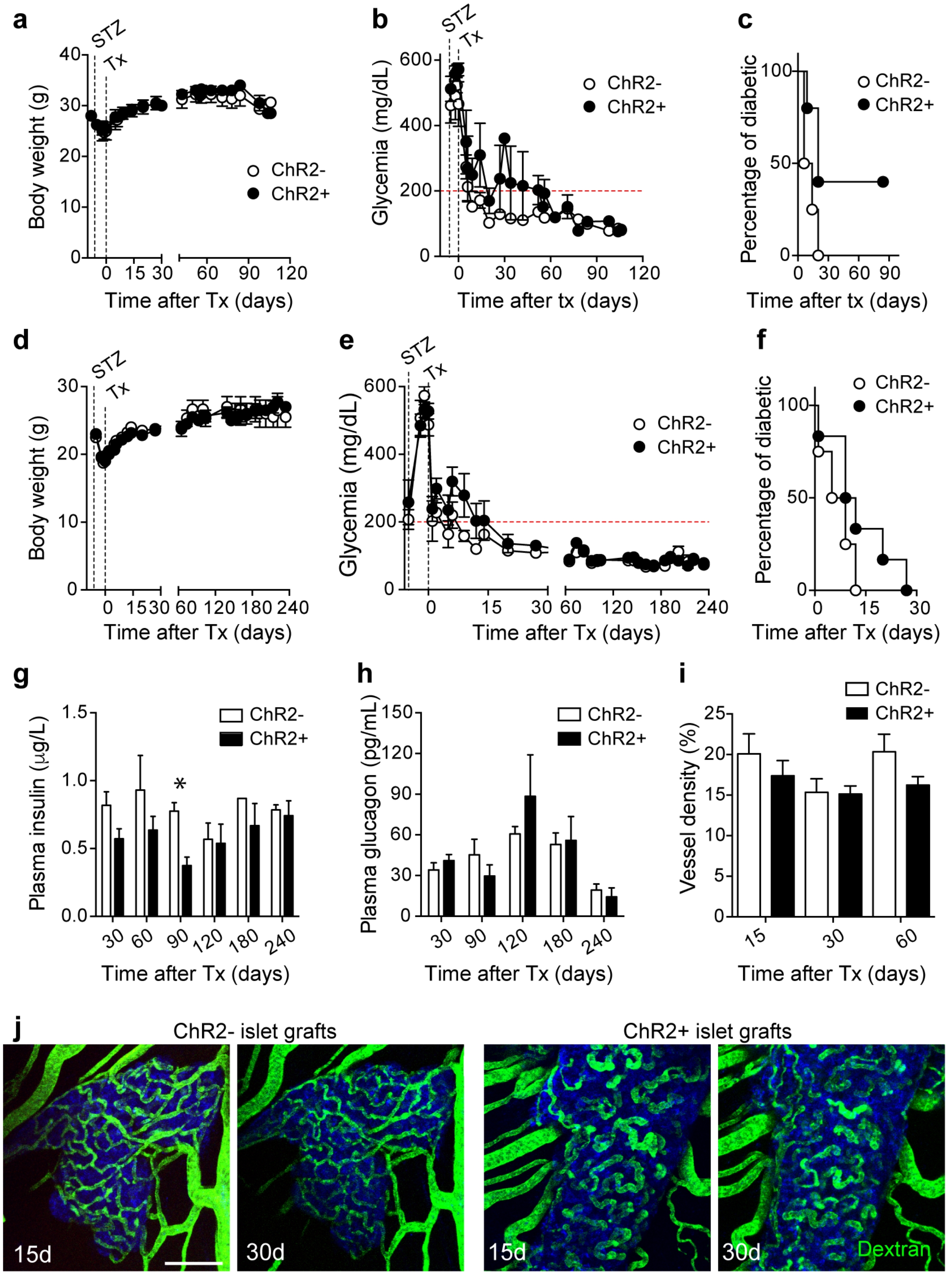


**Supplementary Fig. 2. Transplantation into the anterior chamber of the eye for selective manipulation of islet pericytes.** To assess the effects on glucose metabolism of selective manipulation of islet pericytes and avoid confounding systemic effects, we transplanted islets from mice that express ChR2 in pericytes (ChR2+) into the eyes of male (**a-c;** n=5) and female (**d-f;** n=6) Nude mice (3 months old). As control, we also transplanted islets from mice that do not express ChR2 in pericytes (ChR2-; n=4 male and 4 female recipients). Male and female Nude mice were rendered diabetic (with a STZ injection) before islet transplantation. Islet transplantation reversed hyperglycemia in all the animals that received ChR2- islets (n=8 pooled genders), and in 9 out of 11 mice that received ChR2+ islets (both animals that did not become normoglycemic after transplantation were males; (**c**)). Mice increased their body weight when normoglycemia was achieved suggesting full recovery from STZ treatment. **g,h**, Measurements of non-fasting plasma insulin and glucagon at different time points after transplantation. **i,j**, To monitor the engraftment process, we followed islet revascularization longitudinally by imaging the eye at different time points after transplantation. Animals were injected intravenously with a fluorescent dextran (green). Vessel density in NG2-ChR2+ and NG2-ChR2- grafts at different time points was similar.

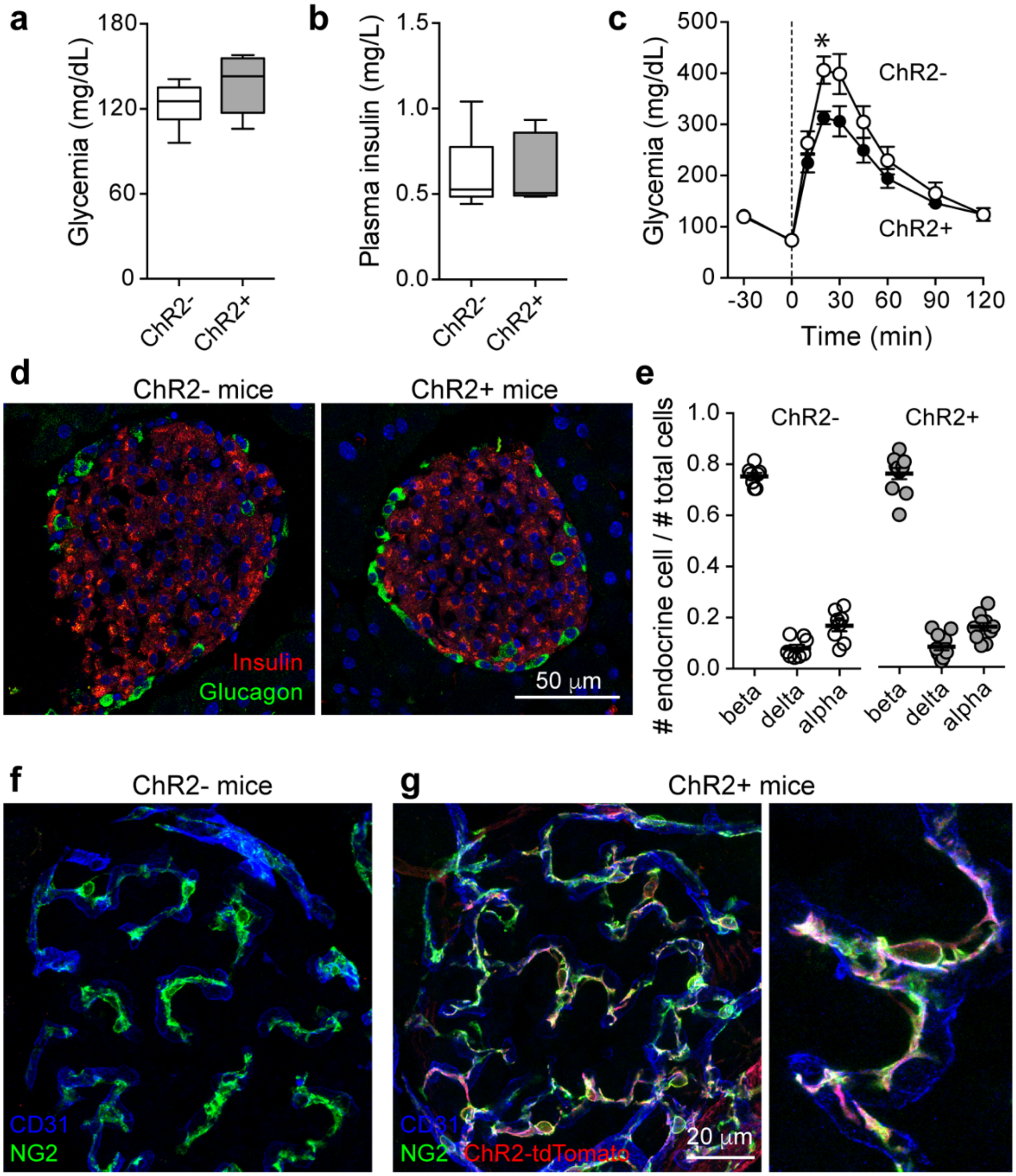


**Supplementary Fig. 3. Transgenic mice for optogenetic manipulation of pericytes.** Pericytes are excitable cells whose contractile activity requires changes in cytosolic calcium levels [3]. To manipulate their activity, we generated mice that express the blue light sensitive ion channel channelrhodopsin2 in pericytes using the Cre/Lox system (see Methods). We generated mice that express ChR2 in pericytes (Cre+; ChR2+) and mice that do not express ChR2 in pericytes (Cre-; ChR2-). Fed glycemia (**a**) and plasma insulin levels (**b**) were not different between donor ChR2+ and ChR2- mice at 3 months of age. **c**, Glucose tolerance was slightly better in ChR2+ mice. **d,e**, We collected pancreases from donor animals and processed for immunohistochemistry. We labeled endocrine cells with antibodies against insulin (beta cells; red), glucagon (alpha cells; green) and somatostatin (delta cells). Islet cytoarchitecture (**d**) and proportion of different endocrine cells (**e**) was not different between ChR2- and ChR2+ donor mice. **f,g**, ChR2 is fused to tdTomato and expressed upon Cre-mediated recombination. NG2-Cre mouse lines direct channels/reporters to smooth muscle cells and pericytes in peripheral tissues such as pancreatic islets. Endothelial cells are shown in blue (CD31) and pericytes are in green (NG2). ChR2 is expressed by pericytes in ChR2+ mice, but not in ChR2- mice.

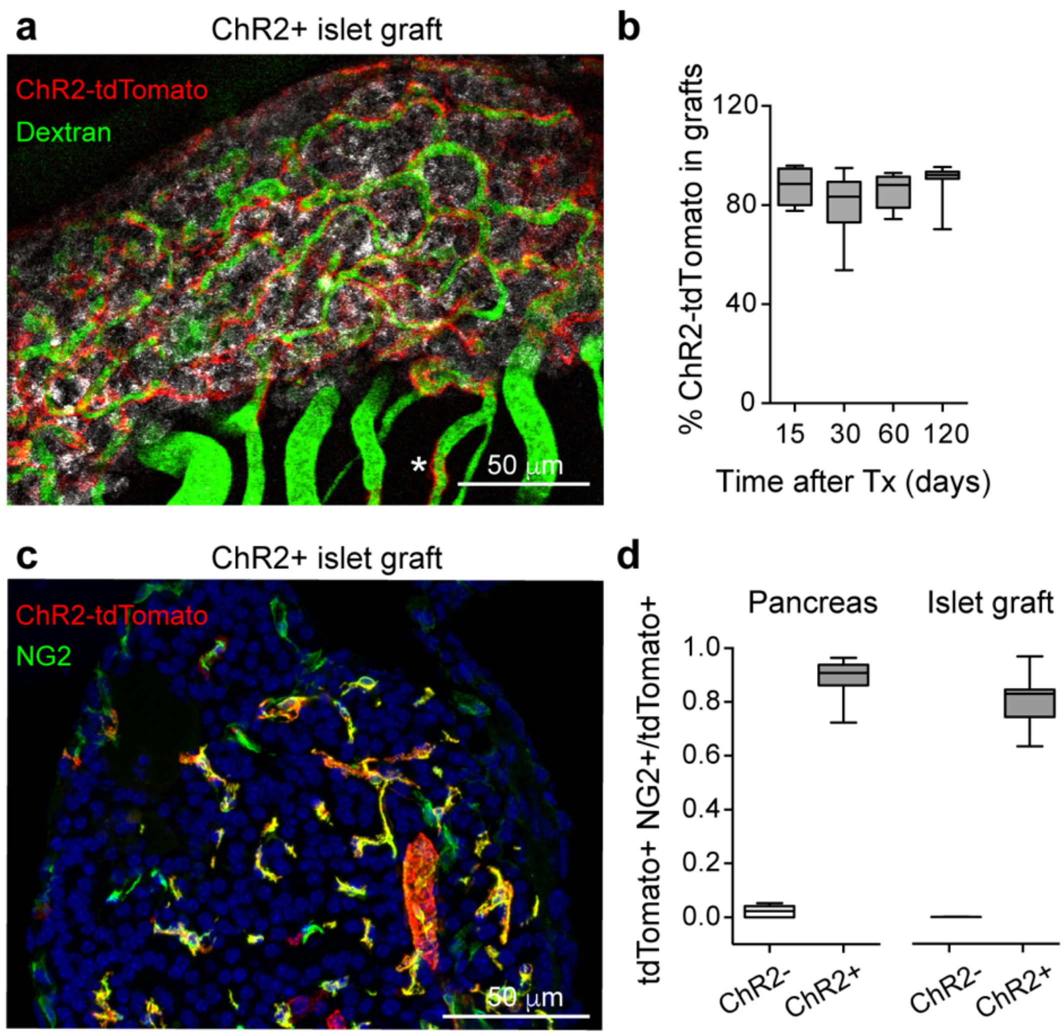


**Supplementary Fig. 4. Islet pericytes retain ChR2 expression after transplantation and remain in grafts**. **a**, ChR2 expressing pericytes (red) and blood vessels (dextran labeled, green) in intraocular islet grafts from NG2-ChR2+ mice can be visualized *in vivo* 3 months after transplantation. * indicates one pericyte that has left the islet graft and is associated with the iris vasculature. **b**, Quantification of the percentage of tdTomato (ChR2) fluorescence that is associated with regions with backscatter (islet grafts). 80-90% of tdTomato expressing cells are in islet grafts. **c**, Eyes with islet grafts were processed for immunohistochemistry and immunostained with an anti-NG2 antibody (green). The majority of tdTomato expressing cells also express NG2. **d**, Mander's coefficient reflecting colocalization of NG2 with tdTomato (ChR2) for islets in the pancreas and grafts in the eye. tdTomato is almost exclusively expressed by NG2 expressing cells both in the pancreas and in the eye.

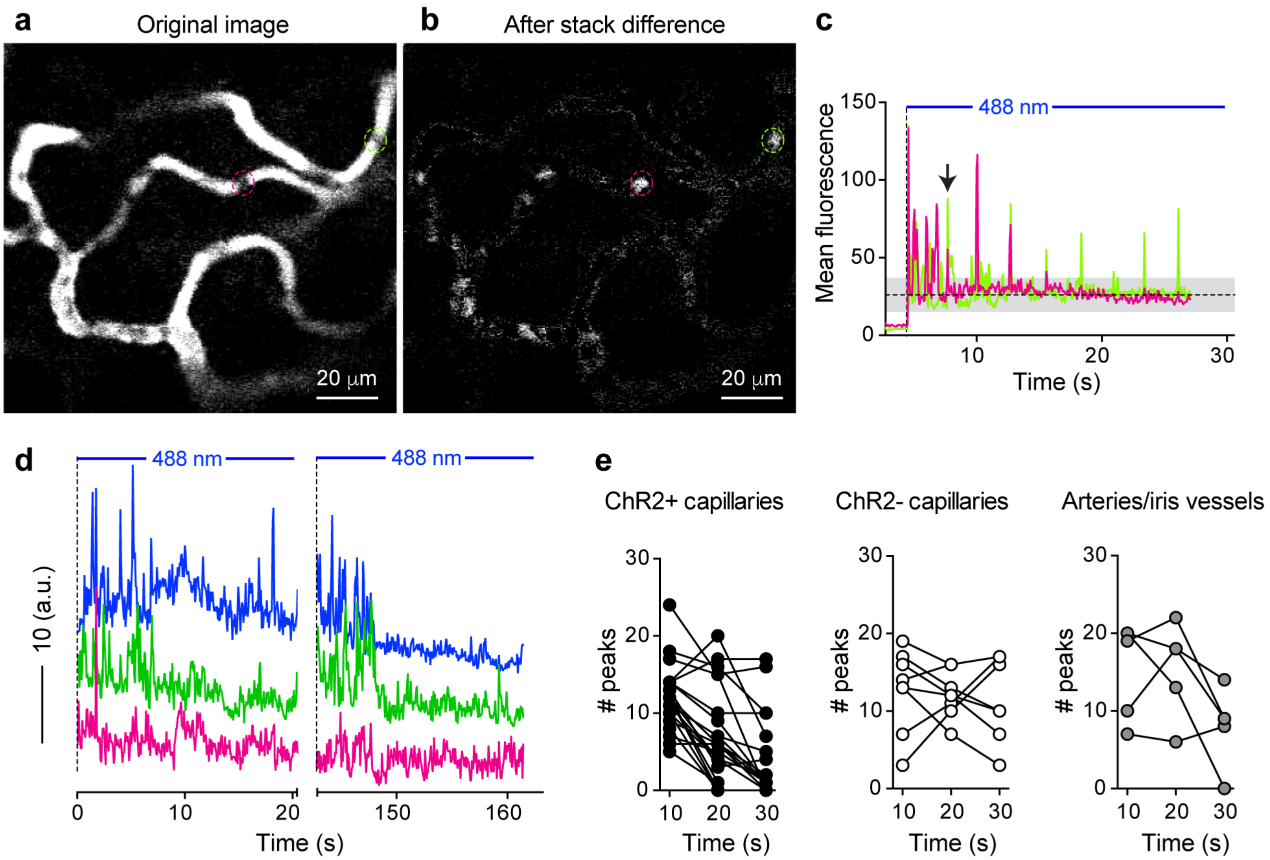


**Supplementary Fig. 5. Pericyte activation with optogenetics decreases blood flow in islet graft capillaries. a**, Confocal image of blood flowing through capillaries in an islet graft from NG2-ChR2+ mice. Image was taken 5s after turning on the 488 nm laser (black arrow in (**c**)). **b**, Corresponding image after applying the "Stack Difference" ImageJ plugin to detect dynamic changes (erythrocyte movement). 2 ROIs were placed on 2 islet capillaries and corresponding changes in fluorescence are shown in **c**. **d**, Changes in blood flow in islet graft capillaries are reversible and reproducible. In a different experiment, the 488 nm laser was turned on twice for 20-30 s. **e**, Quantification of the number of peaks from traces as shown in (**c**) and (**d**) in vessels in grafts from NG2-ChR2+ mice (black symbols), NG2-ChR2- mice (white symbols) or vessels in the iris or at the border of the graft (gray symbols).

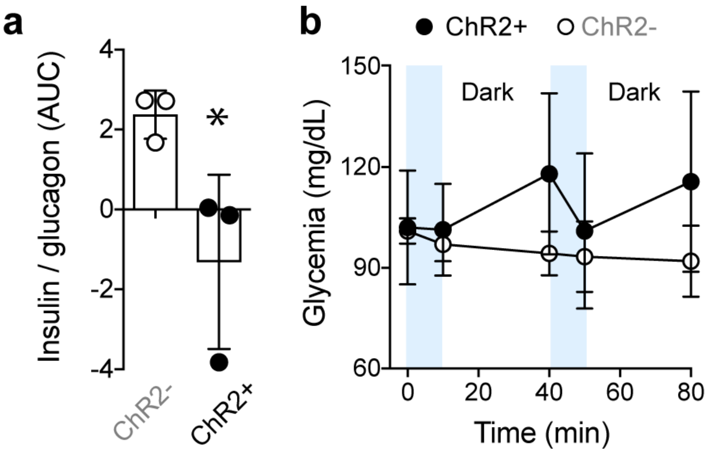


**Supplementary Fig. 6. Pericyte activation with blue light inhibits insulin / glucagon ratio and destabilizes glycemia. a**, Area under the curve of changes in the insulin / glucagon ratio of mice transplanted with NG2-ChR2+ or NG2-ChR2- islets induced by 10 min stimulation with blue light. **b**, Changes in glycemia in transplanted animals placed in the dark and subjected to 10 min interval of blue light stimulation.

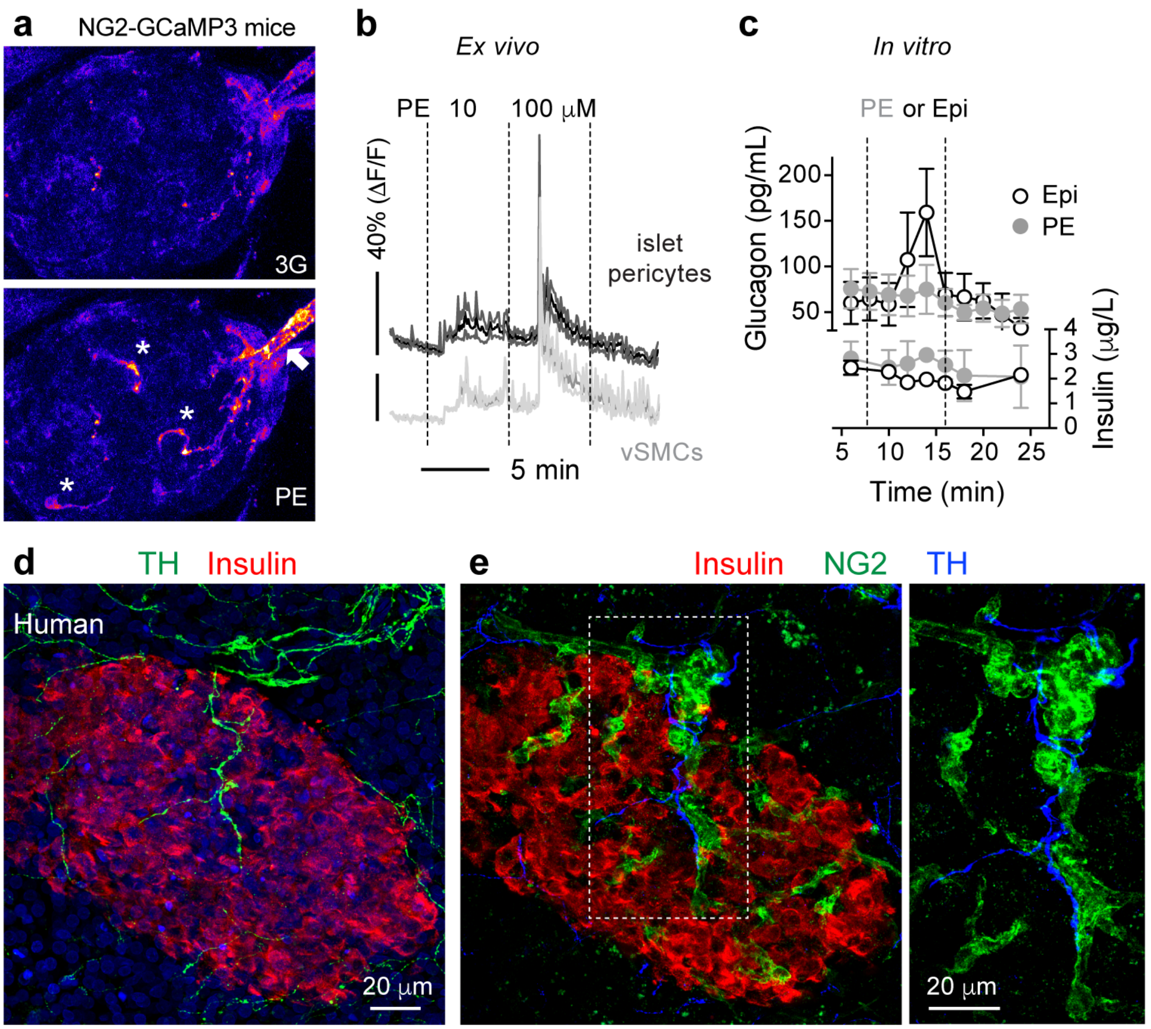


**Supplementary Fig. 7. Islet pericytes are activated by the sympathetic agonist phenylephrine and are the cellular targets of sympathetic axons in the human islet. a**, To assess the effects of the $\alpha$1 adrenergic receptor agonist phenylephrine (PE) in islet pericyte calcium levels ($[Ca^{2+}]i$), we produced living pancreas slices from NG2-GCaMP3 mice. PE increases $[Ca^{2+}]i$ in islet pericytes (indicated with *) and in smooth muscle cells around the islet feeding arteriole (indicated with an arrow). **b**, Traces showing changes in GCaMP3 fluorescence induced by PE (10 and 100 μM), in 3 mM glucose, in regions of interest placed around islet pericytes (dark gray) or smooth muscle cells (light gray). Mean (darker traces) ± SEM (lighter traces) are shown (N = 11 cells/3 mice). **c**, Changes in insulin and glucagon secretion from isolated mouse islets *in vitro* induced by PE (10 μM) or epinephrine (10 μM). PE has no effect on insulin or glucagon secretion *in vitro*, while epinephrine stimulates glucagon secretion and inhibits insulin secretion from isolated mouse islets. **d**, Sympathetic nerves in the human pancreas were visualized with an antibody against tyrosine hydroxylase (TH, green). Insulin-positive beta cells are shown in red. Sympathetic nerves are present in the acinar tissue, at the border of the islet and in the islet parenchyma. **e**, Sympathetic nerves entering the human islet parenchyma follow pericytes (NG2, green) and interact closely with these mural cells (zoomed image). The islet in (**d**) and (**e**) is the same.

### Supplementary Movies

**Supplementary Movie 1. Constriction of islet capillaries induced by optogenetic activation of pericytes.** Movie composed of a series of confocal images taken every 68 msec of ChR2-expressing pericytes (red) in an islet graft in the eye. Endocrine cells are seen in gray (backscattered light). Turning on the 488 nm laser leads to a constriction of the islet vessel indicated with an asterisk. Note that constriction is reversible and, when the laser is turned off, blood flow through that capillary resumes. Movie speed 40 frames per second (fps).

**Supplementary Movie 2. Decrease in blood flowing through islet capillaries induced by optogenetic activation of pericytes.** Movie composed of a series of confocal images taken every 68 msec of islet capillaries (labeled with a dextran, green) covered by ChR2-expressing pericytes (red) in an intraocular graft from a NG2-ChR2+ mice. Endocrine cells are seen in blue (backscattered light). Turning on the 488 nm laser allows us to visualize vessels and leads to a decrease in islet blood flow in some capillaries in the graft. Note that blood flow completely stops in the capillary in the middle of the graft shown with an asterisk. Movie speed 20 frames per second (fps).

**Supplementary Movie 3. Quantifying changes in Decrease in blood flowing through islet capillaries induced by optogenetic activation of pericytes.** Movie composed of a series of confocal images taken every 68 msec of erythrocytes flowing through capillaries in an intraocular islet graft from a NG2-ChR2+ mice. The original movie was processed using the ImageJ plugin “Stack Difference”. Turning on the 488 nm laser decreases blood flow in some vessels in the graft. The capillary located in the middle of the graft corresponds to the vessel shown in Movie S2. Movie speed 20 frames per second (fps).

**Supplementary Movie 4. Decrease in blood flowing through islet capillaries induced by optogenetic activation of pericytes.** Movie composed of a series of confocal images taken every 68 msec of ChR2-expressing pericytes (red) covering islet capillaries (labeled with a dextran, green) in an intraocular graft from NG2-ChR2+ mice. Endocrine cells are seen in blue (backscattered light). Turning the 488 nm laser allows us to visualize the vessels and leads to a decrease in islet blood flow in some vessels in the graft (some responsive vessels are indicated with *). Note that blood flow completely stops in some vessels. Movie speed 40 frames per second (fps).
